# Heterosynaptic cross-talk of pre- and postsynaptic strengths along segments of dendrites

**DOI:** 10.1101/2020.05.05.078543

**Authors:** Rudi Tong, Nigel John Emptage, Yukiko Goda

## Abstract

Dendrites are crucial for integrating incoming synaptic information. Individual dendritic branches are thought to constitute a signal processing unit, yet how neighbouring synapses shape the boundaries of functional dendritic units are not well understood. Here we addressed the cellular basis underlying the organization of the strengths of neighbouring Schaffer collateral-CA1 synapses by optical quantal analysis and spine size measurements. Inducing potentiation at clusters of spines produced NMDA receptor-dependent heterosynaptic plasticity. The direction of postsynaptic strength change showed distance-dependency to the stimulated synapses where proximal synapses predominantly depressed whereas distal synapses potentiated; potentiation and depression were regulated by CaMKII and calcineurin, respectively. By contrast, heterosynaptic presynaptic plasticity was confined to weakening of presynaptic strength of nearby synapses, which required CaMKII and the retrograde messenger nitric oxide. Our findings highlight the parallel engagement of multiple signalling pathways, each with characteristic spatial dynamics in shaping the local pattern of synaptic strengths.

## Introduction

The maintenance of complex neural network dynamics imposes constraints on synaptic strength regulation. Cellular principles underlying neural network function, such as local dendritic integration or the minimization of metabolic cost are thought to give rise to an optimal range of synaptic strength distributions, which in turn, best support network function (Poirazi and Mel 2001; Winnubst and Lohmann 2012; Govindarajan *et al.* 2006; Harris *et al.* 2012). Indeed, synapses display several stereotypical organizational features of their strengths, including an apparent stable distribution across a synapse population that is skewed towards low magnitudes and a non-uniform spatial distribution (Buzsáki and Mizuseki 2014; Barbour *et al.* 2007). How these synaptic patterns emerge from elementary molecular mechanisms is however not fully understood.

Previous studies have reported functional and anatomical similarities between neighbouring synapses (Liang *et al.* 2018; McBride *et al.* 2008; Branco *et al.* 2008; Lee *et al.* 2019; Winnubst and Lohmann 2012; Fu *et al.* 2012; Letellier *et al.* 2019, but see also Jia *et al.* 2010; Chen *et al.* 2011) that were mostly confined to dendritic branches. For instance, pre- and postsynaptic strengths were found to be more similar along short stretches of dendrite than across different branches of dendrites (Branco *et al.* 2008; Letellier *et al.* 2019). Similarly, synapses in close proximity were more likely to be synchronously active (Takahashi *et al.* 2012; Winnubst *et al.* 2015; Kleindienst *et al.* 2011) and functionally correlated (Lee *et al.* 2019; Scholl *et al.* 2017).These synaptic patterns could be either explained by hardwired developmental mechanisms (Druckmann *et al.* 2014; Deguchi *et al.* 2011) or suggest for additional activity-dependent mechanisms that facilitate coordination of synaptic strengths along short dendritic segments. In support of the contribution of activity-dependent mechanisms, pharmacological inhibition of NMDA receptor activity abolished the spatial clustering of temporally correlated inputs (Winnubst *et al.* 2015; Kleindienst *et al.* 2011) and local dendritic activity was shown to adjust pre- and postsynaptic strengths (Branco *et al.* 2008; Letellier *et al.* 2019). Furthermore, functional synaptic clusters were more pronounced after environmental enrichment (Nikonenko *et al.* 2013) and prism rearing in barn owls (McBride *et al.* 2008). Alternatively, these local synaptic patterns may also arise as a consequence of local dendritic spiking (Poirazi *et al.* 2003; Branco and Häusser 2010; Mel 1992) or local homeostatic constraints on resources that promote the normalization of synaptic strengths in a homeostatic manner (Bourne and Harris 2011; Sutton *et al.* 2006; Ju *et al.* 2004).

Collectively, these observations suggest that neighbouring synapses interact with each other and that synaptic plasticity plays a role in implementing this coordination and possibly also competition between synapses. In agreement with such a view of inter-synaptic interactions, a large body of evidence shows that activity-dependent changes of synaptic strength occur at both active (homosynaptic plasticity) and inactive synapses (heterosynaptic plasticity) (Royer and Paré 2003; Schuman and Madison 1994; Scanziani *et al.* 1996; Lynch *et al.* 1977). For instance, changes in synaptic strength at active synapses can either spread to (Engert and Bonhoeffer 1997; Wiegert and Oertner 2013; Hayama *et al.* 2013; Letellier *et al.* 2019) or induce a compensatory response at adjacent synapses (Bian *et al.* 2015; Oh *et al.* 2015; El-Boustani *et al.* 2018; Lee *et al.* 2013) or affect metaplastic properties to bias future modifications (Harvey and Svoboda 2007; Harvey *et al.* 2008; Fonseca *et al.* 2004; Govindarajan *et al.* 2011). A better understanding of when, where, and how neighbouring synapses interact will help provide insights into the emergence of higher level organizational principles of synapses such as the minimal computational unit and how its bound is set (Govindarajan *et al.* 2006).

Although a variety of phenomena and mechanisms have been reported for synaptic interactions, a common framework to explain the spatial coordination of synaptic strengths has been difficult to formulate (Kruijssen and Wierenga 2019). For instance, it has been suggested that low-level organizational patterns might emerge from the unique spatiotemporal properties of signalling molecules that interact during synaptic plasticity (Nishiyama and Yasuda 2015). However, the interplay of parallel plasticity pathways during synaptic plasticity has been difficult to observe and dissociate experimentally. In addition, even though it is generally accepted that synaptic plasticity occurs at both pre- and postsynaptic loci (Bliss and Collingridge 1993) and whilst heterosynaptic signalling between postsynaptic terminals along dendrites has been explored, little to nothing is known about heterosynaptic plasticity at the presynaptic terminal. This can mainly be attributed to the technical difficulties of measuring presynaptic strength, especially in more intact systems such as brain slices or *in vivo* (O’Rourke *et al.* 2012; Burette *et al.* 2015). However, investigating synaptic plasticity at both loci is particularly important because pre- and postsynaptic terminals might represent functionally distinct and independently regulated compartments (Padamsey *et al.* 2017; Costa *et al.* 2017; Xu *et al.* 2013; Sáez and Friedlander 2009; Letellier *et al.* 2019) and are therefore likely to also differ in their spatial regulation with consequences on the property of information transfer (Costa *et al.* 2017).

Here, we have addressed the cellular basis underlying the organization of neighbouring synaptic strengths at the Schaffer collateral inputs to CA1 pyramidal neurones. Using a combination of optical quantal analysis (Padamsey *et al.* 2019) and spine size measurements at individual synapses, we elicited potentiation at clusters of spines and assessed pre- and postsynaptic strength changes at nearby synapses. Our findings reveal a spread of plasticity along CA1 pyramidal neurone dendrite segments with a pattern distinct between the pre- and the postsynaptic sides, and our observations point to the engagement of independent signalling pathways that operate in parallel.

## Results

### Local structural potentiation of groups of spines induces bi-directional postsynaptic heterosynaptic plasticity

We first investigated the synaptic cross-talk that might contribute to the formation of stable functional clusters. We hypothesized that the local coordination of synaptic strength is implemented by a tight balance of cooperation and competition between synapses. This predicts that experimentally imposing strong coordinated activity locally at a subset of synapses will induce a reorganization of synaptic strengths at all nearby synapses due to underlying competitive processes. We therefore emulated the activation of groups of spatially clustered synapses along segments of dendrites using 2-photon glutamate uncaging. Groups of synapses were stimulated quasi-synchronously, and this stimulation was additionally paired with cell-wide depolarisation of the postsynaptic neurone to promote dendritic signalling events for the expression of synaptic plasticity (**Fig. 1A**). In order to monitor postsynaptic strength over the course of the experiment, we estimated changes in spine volume, a parameter known to tightly correlate with the number of functional postsynaptic AMPA receptors (Matsuzaki *et al.* 2001; Matsuzaki *et al.* 2004; Smith *et al.* 2003; Béïque *et al.* 2006). As a readout of presynaptic strength, we used optical quantal analysis to determine release probability (Pr) at selected synapses (Padamsey *et al.* 2019). **Figure 1** summarizes the experimental setup. We first bolus-loaded (45-60 s) CA1 pyramidal neurones in organotypic hippocampal slices via the patch pipette with the Ca^2+^ indicator OGB-1 (1 mM) and a fluorescent dye (AF594, 0.5 mM) to visualize fine sub-cellular structures (**Fig. 1A,B**). Next, we placed a stimulation electrode close to a dendritic segment of interest (**Fig. 1B**) and measured presynaptic strength of select synapses using optical quantal analysis (**Fig. 1C**). For glutamate uncaging, caged glutamate (10 mM MNI-glutamate) was locally perfused and we adjusted the laser power of the uncaging laser to match the magnitude and temporal dynamics of Ca^2+^ responses evoked by electrical stimulation (**Fig.1D**). To elicit LTP, five to seven adjacent dendritic spines were targeted by quasi-synchronous glutamate photolysis (30x, 0.5 Hz) paired with depolarisation of the patch-clamped cell to 0 mV (**Fig. 1E**). We re-examined presynaptic strength after 30 min by repeating optical quantal analysis; spine structure was monitored for up to 45 min post-induction. We term LTP induced by this form of clustered stimulation as”cl-LTP”.

**Fig. 1:**
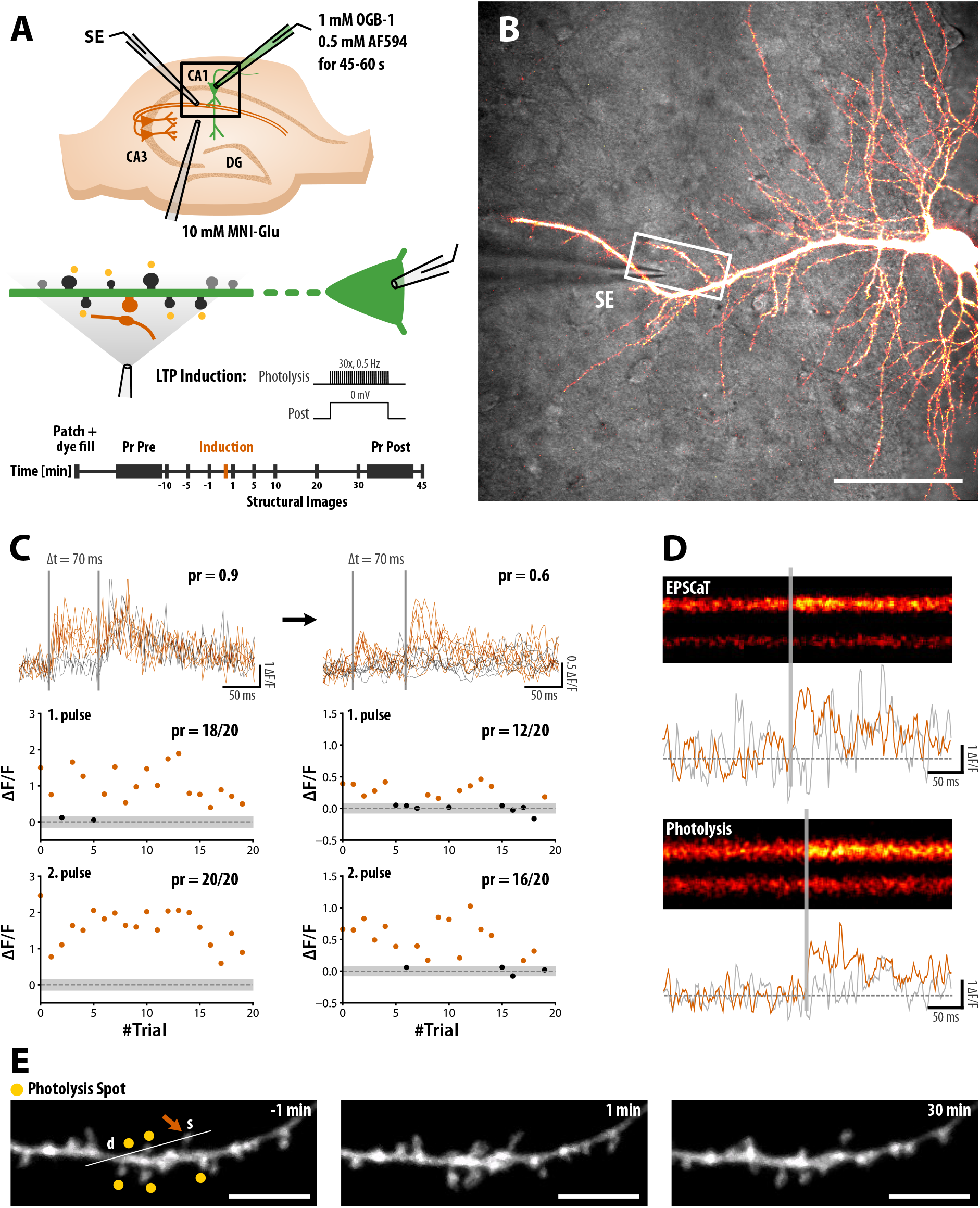
Experimental setup to study pre- and postsynaptic heterosynaptic plasticity. (A) Schematic of the experimental setup (top), cl-LTP induction protocol (middle), and timeline (bottom). (B) Example neurone filled with 1 mM OGB-1 and 0.5 mM AF594 for 1 min. The white box illustrates the region shown in (E). SE: Stimulation electrode. Scale bar: 20 μm. (C) Example of Pr measurements via optical quantal analysis before (left) and after (right) cl-LTP induction. Raw fluorescence traces of recordings (top) sorted into two groups showing either successful release (red) or unsuccessful release (grey) to the first AP. The plots below show the peak intensity of the response to the first (middle) and second pulse (bottom). The thick grey bar corresponds to two standard deviations of the baseline, which was used as detection criteria for automatic analysis. Facilitation of the second pulse is clearly evident as an increase in Pr. (D) Glutamate photolysis was adjusted to elicit Ca^2+^ transients similar in strength compared to EPSCaTs. The grey trace corresponds to the nearby dendrite. (E) cl-LTP was induced via quasi-synchronous photolysis of glutamate at 5-7 spines paired with postsynaptic depolarisation, 30x at 0.5 Hz. An increase in spine size can be seen immediately after photolysis. For optical quantal analysis, line scans were taken through the spine of interest (s) and the dendritic branch (d). See Methods for details. Scale bar: 10 μm.

Clustered stimulation led to structural enlargement of spine head volumes of a large fraction of stimulated spines, which was stable for at least 45 min (cl-LTP; **Fig. 2Ci,** 59.5 ± 4.1 % of spines with > 20 % increase in spine size at 30-45 min). The average increase in spine size after 30-45 min was ΔV_30-45min_ = 38.7 ± 4.5 % of baseline (N = 17 experiments). Neighbouring unstimulated spines also transiently increased in size shortly after cl-LTP induction (**Fig. 2Ci,** ΔV_10min_ = 10.8 ± 4.0 %) before slowly decaying back to baseline by 45 min (ΔV_30-45min_ = −1.1 ± 2.2 %). When we prevented postsynaptic depolarisation by clamping the cell at −70 mV during clustered stimulation, stimulated spines did not increase in size, neither transiently nor over the long-term (**Fig. 2Di**, ΔV_30-45min_ = −4.9 ± 5.9%, N = 9). Without postsynaptic depolarisation, neighbouring unstimulated spines also remained unchanged (**Fig. 2Di**, ΔV_10min_ = −2.8 ± 4.2 %; ΔV_30-45min_ = −3.1 ± 2.9 %). Local application of AP5 (500 μM, locally perfused) also abolished both homo- and heterosynaptic spine structural changes (**Fig. S1Ai**).

**Fig. 2:**
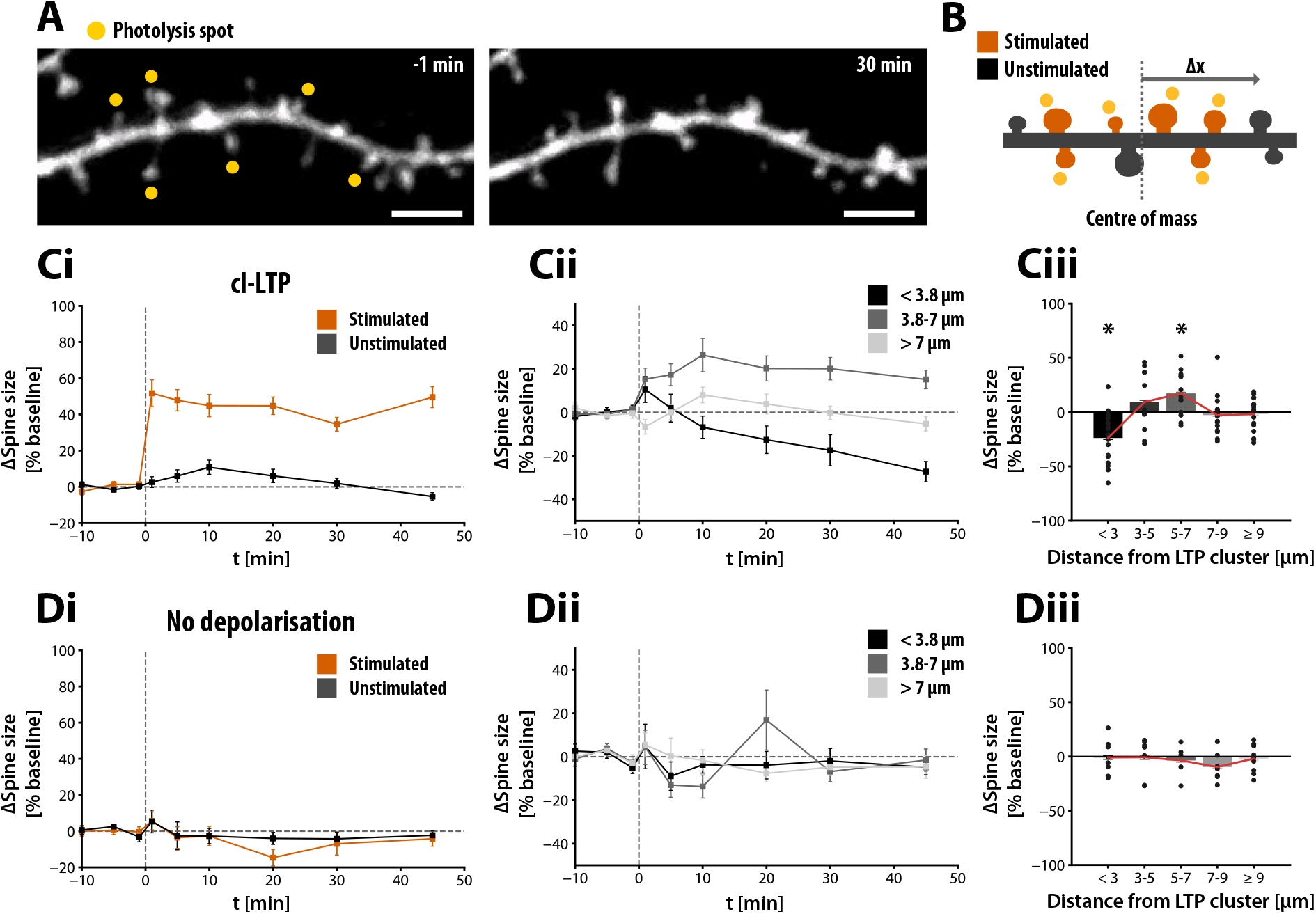
Induction of structural plasticity of groups of spines leads to heterosynaptic bidirectional changes in spine size. (A) Example of cl-LTP induction at six spines via photolysis of glutamate. Long-term spine structural changes were assessed for 30-45 min. Scale bar: 5 μm. (B) The distance metric used for analysis of the spine cluster was defined as the distance along the dendritic branch to the centre of mass of the stimulated cluster (Δx). (Ci) Spine size changes of stimulated and unstimulated spines. cl-LTP was induced at t = 0 min. Stimulated spines showed robust structural LTP (ΔV_30-45min_ = 38.7 ± 4.5 %, N = 17), whereas unstimulated spines enlarged transiently (ΔV_10min_ = 10.8 ± 4.0 %, ΔV_30-45min_ = −1.1 ± 2.2 %). (Cii) Spine size changes of unstimulated spines, grouped by distance (< 3.8 μm: ΔV_30-45min_ = −23.0 ± 5.3%; 3.8-7 μm: ΔV_30-45min_ = 18.1 ±4.0%; > 7 μm: ΔV_30-45min_ = −2.3 ±2.9%, see also **Figure S2**). Spines located close to the stimulated cluster decreased in size. More distal spines, however, showed significant increase in size. (Ciii) Persistent spine size change at 30-45 min depends on the distance to the stimulated cluster. Spines belonging to within 3 μm and 5-7 μm groups were significantly different from control experiments summarized in (Diii) (Mann-Whitney U test). The polarity change associated with distance-dependency of the mean spine size change is highlighted in red. (D) Same as C but postsynaptic depolarisation was prevented by holding the neurone at −70 mV during cl-LTP induction. Neither stimulated nor unstimulated spines showed substantial structural changes. Data points are shown as mean ± SEM.

Spatial proximity plays an important role in heterosynaptic plasticity, since spatially delimited diffusion of signalling molecules between synapses is thought to mediate the local coordination of synaptic strengths (Nishiyama and Yasuda 2015). We therefore grouped unstimulated spines based on their distance along the dendrite to the centre of mass of the stimulated cluster (**Fig. 2B**, see Methods). Unstimulated spines within 7 μm of the stimulated cluster showed significant bidirectional structural changes (**Fig. S2**). The distance-dependence of the bidirectional changes was most apparent when comparing spines within and spines beyond 3.8 μm of the stimulated cluster (**Fig. S2**). We found that spines located within 3.8 μm showed a consistent decrease in size, which lasted for at least 45 min (**Fig. 2Cii**, ΔV_30-45min_ = −23.0 ± 5.3 %). By contrast, spines at a distance of 3.8-7 μm exhibited weak but significant potentiation, which lasted for the duration of the experiment (ΔV_30-45min_ = 18.1 ± 4.0 %). Spines located further away showed a slow but transient increase in size, reaching a maximum at 10 min after stimulation and decaying back to baseline by the end of the experiment (ΔV_30-45min_ = −2.3 ± 2.9 %). **Fig. 2Ciii** summarizes the distance-dependent bidirectional regulation of postsynaptic strength, which is absent without postsynaptic depolarisation (**Fig. 2Diii**, cl-LTP vs. no depolarisation: p < 0.05 for spines located < 3 μm and 5-7 μm, Mann-Whitney U test) and when NMDAR signalling was blocked with AP5 (**Fig. S1**).

### Local structural potentiation of groups of spines induces presynaptic heterosynaptic weakening

Next, we examined presynaptic changes following cl-LTP. Due to the comparatively low throughput nature of optical quantal analysis in which Pr measurement in general was feasible for only a single synapse per experiment, we focused mainly on synapses located close to the stimulated cluster (synapse located < 4 μm to the stimulated cluster in 13/17 experiments). cl-LTP induction resulted in a decrease in Pr of synapses located close to the stimulated cluster (**Fig. 3C**, ΔPr = −0.27 ± 0.05, N = 13, all spines located within 4 μm), whereas Pr of synapses located more distally did not change (**Fig. 3C** triangles, ΔPr = −0.02 ± 0.06, N = 4, all spines located > 4 μm). Given that the activation of presynaptic NMDA receptors can lead to the expression of presynaptic LTD (Padamsey *et al.* 2017), the large amounts of glutamate released during photolysis could have resulted in local accumulation and spill-over of glutamate to cause a decrease in Pr at neighbouring unstimulated synapses. However, holding the neurone at −70 mV during cl-LTP induction abolished presynaptic changes (ΔPr = −0.02 ± 0.06, N = 11, p < 0.01, Kruskal-Wallis H-test with *post hoc* Dunn’s test, **Fig. 3D,** “no depolarisation”). Thus glutamate uncaging by itself was not sufficient to induce heterosynaptic LTD of Pr, and concurrent postsynaptic depolarisation was required. Notably, postsynaptic depolarisation alone is known to influence Pr (Branco *et al.* 2008;Volgushev *et al.* 1997).Therefore, to control for this possibility, we repeated the experiment without glutamate photolysis by either locally perfusing aCSF without caged glutamate or by omitting the photolysis pulse while applying postsynaptic depolarisation. Under these conditions, presynaptic weakening was not observed (**Fig. 3D** “no photolysis”, ΔPr =-0.01 ± 0.03, N = 18, p < 0.01, Kruskal-Wallis H-test with *post hoc* Dunn’s test). Presynaptic heterosynaptic LTD was also abolished by the local application of AP5 during the induction of cl-LTP (**Fig. S1C**). Thus, presynaptic heterosynaptic long-term depression (hetLTD) requires both postsynaptic depolarisation and synaptic stimulation of nearby synapses.

**Fig. 3:**
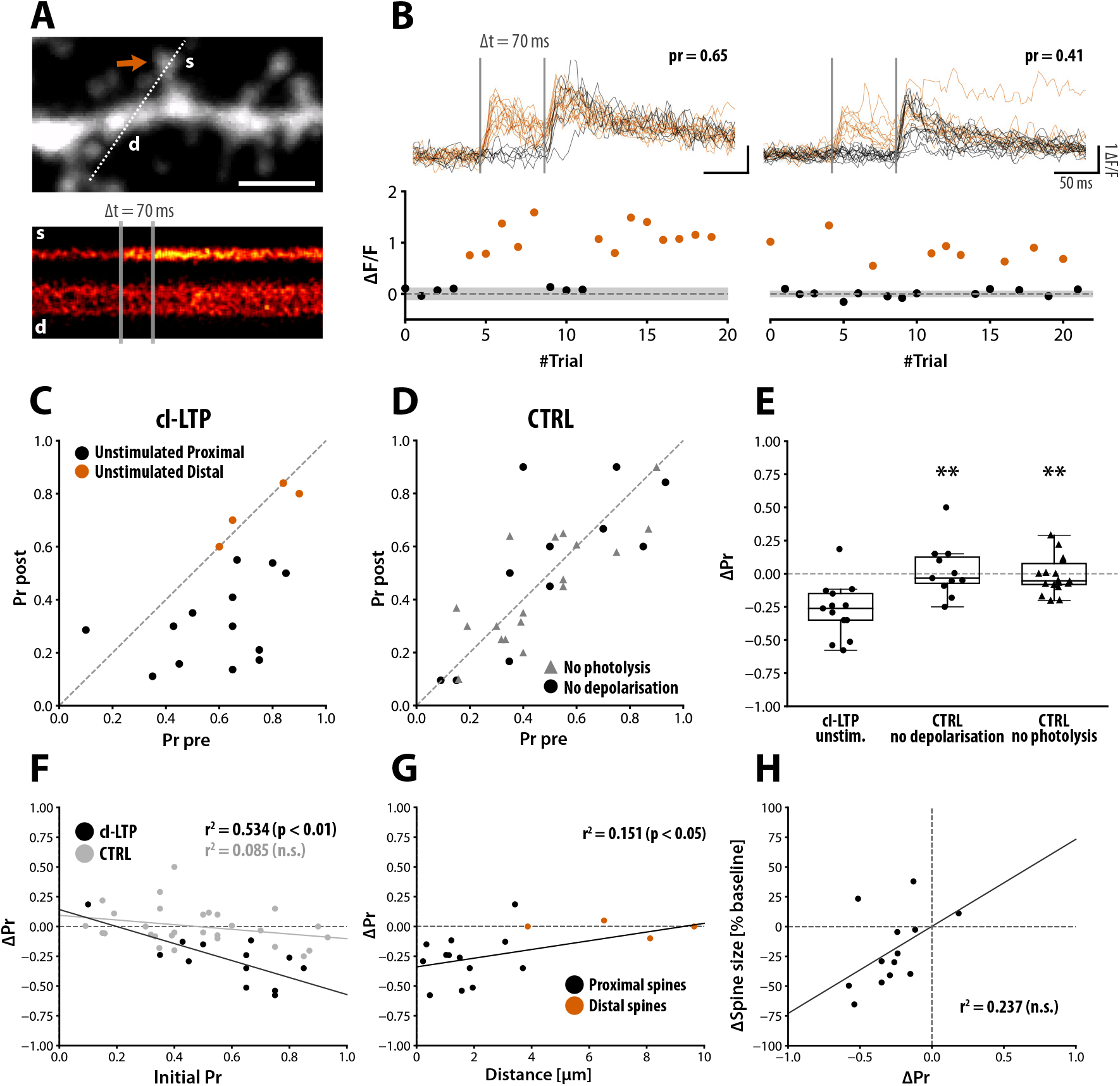
Induction of structural plasticity of groups of spines leads to heterosynaptic presynaptic weakening. (A) Pr was measured using optical quantal analysis. Top: Line scans were taken through the spine of interest (red arrow, s) and the dendrite (d) while the synapse was stimulated. Bottom: Glutamate release was detected as Ca^2+^transients in the spine head (ESPCaTs). (B) Top: raw Ca^2+^ transients show all-or-none response profile to repeated stimulation, before (left) and after (right) the induction of cl-LTP. Bottom: Peak amplitude of the EPSCaT. The grey bar represents two standard deviations from the baseline. An increase in release failures is observed 30 min after cl-LTP induction (right panel). (C) Pr 30 min after cl-LTP induction (post) versus baseline Pr (pre). Pr decreased for synapses located close to the stimulated cluster (< 4 μm, “Unstimulated Proximal”) but not in distal synapses (> 4 μm,”Unstimulated Distal”). (D) No Pr change was observed when omitting glutamate photolysis (“No photolysis’, triangles) or postsynaptic depolarisation (“No depolarisation’, circles). (E) Summary of Pr changes (cl-LTP unstim: ΔPr = −0.27 ± 0.05, N = 13, all synapses within 4 μm of the stimulated cluster; CTRL no depol.: ΔPr = −0.02 ± 0.06, N = 11; CTRL no photolysis: ΔPr = −0.01 ± 0.03, N = 18). Significance is shown for comparisons with “cl-LTP unstim” (Kruskal-Wallis H-test with *post hoc* Dunn’s test). (F) The change in Pr as function of the initial Pr. (G) Pr changes relative to distance to the stimulated cluster. (H) Correlation between pre- and postsynaptic heterosynaptic changes.

The decrease in Pr at synapses neighbouring the stimulated spines was correlated with the initial Pr (**Fig. 3F, “black circles”**), which we did not observe for control conditions (**Fig. 3F, “grey circles”**). This dependence of the extent hetLTD of Pr on the initial Pr, which in itself was highly heterogeneous (mean initial Pr = 0.58 with standard deviation of 0.21) could explain the variance of the data for the change in Pr as a function of distance. Specifically, the magnitude of presynaptic changes was not strongly related to the distance to the stimulated cluster (**Fig. 3G**). Instead, the observation of presynaptic hetLTD was limited to synapses located within ~ 4 μm of the stimulated cluster, unlike the magnitude of spine size changes which were correlated with the distance to the stimulated cluster for up to ~ 7 μm. Moreover, we did not observe a significant correlation of pre- and postsynaptic changes (Pearson r = 0.487, N = 13, p = 0.09, **Fig. 3H**).

Altogether, strong stimulation of groups of synapses led to a restructuring of synaptic strengths along the dendrite. The postsynaptic strength of unstimulated synapses changed bi-directionally, which was dependent on the distance to the stimulated synapses. In contrast, the presynaptic terminal weakened only at synapses close to the stimulated cluster. The different patterns of heterosynaptic pre- and postsynaptic changes and the lack of correlation between pre- and postsynaptic changes at individual synapses suggest the parallel engagement of independent signalling pathways.

### Presynaptic hetLTD requires activation of CaMKII, whereas postsynaptic changes are driven by parallel signalling pathways involving calcineurin and CaMKII

Next, we sought to investigate the underlying molecular pathways involved in the heterosynaptic cross-talk. The spatial reorganization of synaptic strengths might reflect the biochemical computations driven by the interaction of signalling pathways (Nishiyama and Yasuda 2015). There is strong evidence that heterosynaptic postsynaptic weakening requires the activation and diffusion of the phosphatase calcineurin (Oh *et al.* 2015; Hayama *et al.* 2013). In contrast, the activation of the kinase CaMKII and its downstream targets including the small GTPases was reported to facilitate LTP at nearby synapses (Harvey and Svoboda 2007; Harvey *et al.* 2008; Hedrick *et al.* 2016).Therefore, calcineurin and CaMKII are potential candidates for plasticity factors mediating the depression and potentiation, respectively. Our observation of a bi-directional regulation of postsynaptic strength could be explained by the orchestrated spatial diffusion of these two plasticity factors, with CaMKII causing potentiation and calcineurin causing depression. We thus predicted that 1) selective inhibition of calcineurin should abolish the postsynaptic weakening of proximal synapses without affecting the potentiation at more distal synapses and 2) inhibition of CaMKII should lead to the opposite effect. To test this, we repeated the experiment under the pharmacological blockade of either calcineurin or CaMKII.

The vehicle control recapitulated our previous results (**Fig. 2**) with the exception of a slight shift in the spatial bounds that best explained the bi-directional heterosynaptic changes (**Fig. 4A**, < 2.8 μm: ΔV_30-45min_ = −15.5 ± 4.7 %; 2.8-6.5 μm: ΔV_30-45min_ = 11.4 ± 5.7 %; > 6.5 μm: 5.9 ± 3.1 %; see also **Fig. S2**), which could have been attributed to culture variability. In the presence of bath applied 10 μM KN62, an inhibitor of CaMKII, the cl-LTP of stimulated spines was blocked (**Fig. 4B,** ΔV_30-45min_ = 6.4 ± 9.2 %, N = 10, p < 0.01), consistent with previous reports (Matsuzaki *et al.* 2004). In the presence of KN62, unstimulated spines located close to the stimulated spines decreased in size as observed above, however, similarly to the stimulated spines, the enlargement of spines located more distally was completely abolished (**Fig. 4Bii**, < 2.8 μm: ΔV_30-45min_ = −10.6 ± 8.2 %; 2.8-6.5 μm: ΔV_30-45min_ = −6.9 ± 5.6 %; > 6.5 μm:ΔV_30-45min_ = −3.1 ±3.2 %). In contrast, inhibition of calcineurin by bath application of 2 μM FK506 completely abolished heterosynaptic postsynaptic weakening of proximal spines (**Fig. 4C**). Instead, upon blocking calcineurin activity, we observed potentiation of both proximal and distal spines (< 2.8 μm:ΔV_30-45min_ = 24.7 ±8.8%; 2.8-6.5 μm:ΔV_30-45min_ = 17.7 ± 8.3 %; > 6.5 μm: ΔV_30-45min_ =6.6 ± 4.9 %). FK506 did not affect the magnitude of cl-LTP at stimulated spines (ΔV_30-45min_ = 40.4 ± 4.3 %, N = 8). Asummary ofthedata is shown in **Fig. 4D**. Inhibition of CaMKII but not calcineurin abolished also presynaptic heterosynaptic weakening (**Fig. 4E**).

**Fig. 4:**
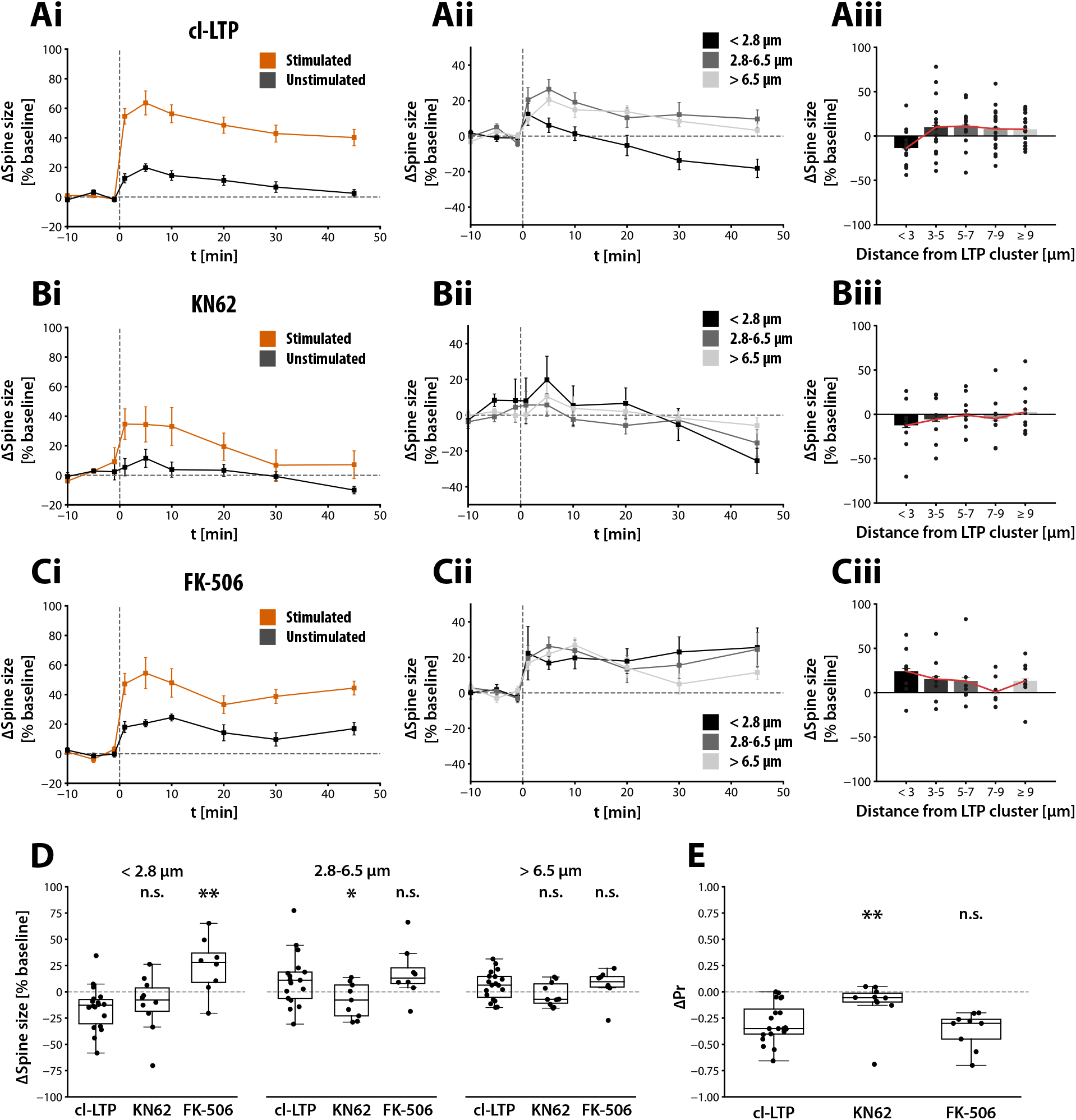
CaMKII is required for postsynaptic heterosynaptic LTP whereas calcineurin is required for postsynaptic heterosynaptic LTD. (Ai) Spine size changes of stimulated (orange) and unstimulated (grey) spines. Spine enlargement was robust and persistent at stimulated spines (ΔV_30-45min_ = 42.5 ± 5.1 %, N = 19) and transient at unstimulated spines (ΔV_10min_ = 14.6 ± 3.3 %). (Aii) Spine size changes of unstimulated spines, grouped by distance (< 2.8 μm: ΔV_30-45min_ = −15.5 ±4.7 %; 2.8-6.5 μm: ΔV_30-45min_ = 11.4 ± 5.7 %; > 6.5 μm: 5.9 ± 3.1 %). (Aiii) Spine size changes of unstimulated spine, grouped in 2 μm bins. (Bi-iii) Effects of bath applied CaMKII inhibitor KN62 (10 μM) on spine size changes. The expression of homo- and heterosynaptic LTP but not LTD was abolished (N = 7, stimulated: ΔV_30-45min_ = 6.4 ± 9.2 %; < 2.8 μm: ΔV_30-45min_ = −10.6 ± 8.2%; 2.8-6.5 μm: ΔV_30-45min_ = −6.9 ±5.6 %; > 6.5 μm: ΔV_30-45min_ = −3.1 ± 3.2%). (Ci-iii) Effects of bath applied calcineurin inhibitor FK-506 (2 μM). Heterosynaptic shrinkage of proximal spines was completely abolished and spine enlargement was observed instead (N = 8, < 2.8 μm: ΔV_30-45min_ = 24.7 ± 8.8 %; 2.8-6.5 μm: ΔV_30-45min_ = 17.7 ± 8.3 %; > 6.5 μm: ΔV_30-45min_ = 6.6 ± 4.9 %). Homosynaptic LTP remained unchanged (ΔV_30-45min_ = 40.4 ± 4.3 %, N = 8). Error bars represent SEM. (D) Comparison of heterosynaptic spine size changes for proximal (< 2.8 μm), distal (2.8-6.5 μm), and more distant spines (> 6.5 μm). Statistical comparison includes L-NAME experiments shown in Fig. 5 for the multiple comparison. (E) Comparison of heterosynaptic Pr changes for spines located within 4 μm of the stimulated spine cluster (Kruskal-Wallis H-test with *post hoc* Dunn’s test). Significance indicates comparison with cl-LTP group. Data points are shown as mean ± SEM.

Taken together, these experiments demonstrate a clear pharmacological dissociation of heterosynaptic postsynaptic potentiation and depression. Consistent with our hypothesis, our observations suggest that postsynaptic heterosynaptic plasticity is regulated by at least two opposing and likely diffusible factors, which depend on calcineurin and CaMKII activity. Moreover, selective block of presynaptic but not postsynaptic weakening by KN62 points also to a dissociation between pre- and postsynaptic plasticity. Notably, upon inhibiting calcineurin, presynaptic weakening was accompanied by postsynaptic potentiation. Presynaptic heterosynaptic LTD is therefore not likely caused by a matching between pre- and postsynaptic strengths (see e.g.Tang *et al.* 2016; Meyer *et al.* 2014; Bayazitov *et al.* 2007; Kay *et al.* 2011) but is subject to regulation by an independent signalling pathway that acts in parallel to the postsynaptic regulatory mechanism.

### Nitric oxide is the retrograde messenger for presynaptic heterosynaptic LTD

The heterosynaptic presynaptic weakening in our experimental paradigm necessitated some form of retrograde signalling as the effects of stimulation targeted to the postsynaptic spines were prevented in the absence of depolarisation of the postsynaptic neurone. The findings so far suggested that the retrograde messenger was likely produced downstream of NMDA receptor activation, involved CaMKII signalling either upstream or downstream, and was expected to rapidly diffuse over short distances from the stimulated synapse cluster. Any one of a range of known candidate molecules fulfills these requirements. In particular, strong local dendritic depolarisation can trigger the production and release of nitric oxide (NO) (Padamsey *et al.* 2017), a highly diffusible signalling molecule. Furthermore, NO signalling is known to interact with NMDA receptor and CaMKII signalling (Hardingham *et al.* 2013). NO is also implicated in (homosynaptic) presynaptic LTP (Padamsey *et al.* 2017) and LTD (Stanton *et al.* 2003). We therefore hypothesized that strong local activity triggers the production and release of NO, which due to its gaseous nature, diffuses over a confined distance to neighbouring presynaptic terminals to elicit presynaptic weakening. We tested this by applying the NO synthase inhibitor L-NAME.

Bath application of 100 μM L-NAME completely abolished heterosynaptic presynaptic LTD (**Fig. 5B,D**, ΔPr = −0.02 ± 0.06, N = 8, p < 0.01). By contrast, L-NAME did not affect cl-LTP induction and expression at stimulated spines (ΔV_30-45min_ = 53.6 ± 9.5 %, N = 8), the transient increase in spine size immediately after induction (ΔV_10min_ = 11.7 ± 1.4 %), or the distance-dependent bi-directional heterosynaptic regulation of spine size (**Fig. 5A**). In fact, the fraction of stimulated spines expressing stable structural LTP was significantly larger when NO production was inhibited (77.4 ± 4.5 % of spines with ΔV_30-45min_ > 20 %, p < 0.05 vs. vehicle).

**Fig. 5:**
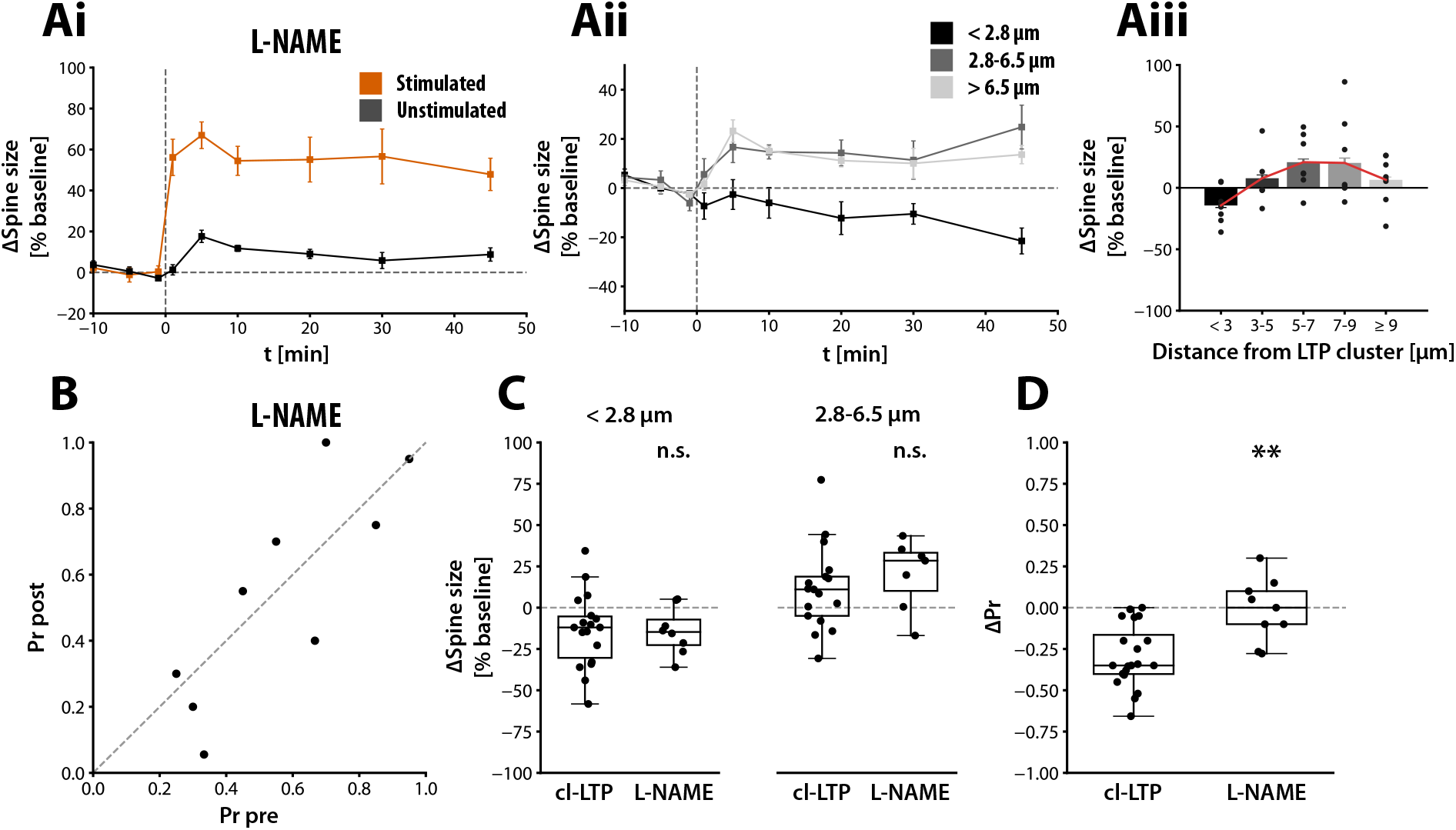
Nitric oxide is the retrograde messenger for heterosynaptic presynaptic weakening. Spine size changes in the presence of bath applied NO synthase inhibitor L-NAME (100 μM). (Ai) Spine size changes of stimulated (ΔV_30-45min_ = 53.6 ± 9.5 %, N = 8) and unstimulated spines (ΔV_10min_ = 11.7 ± 1.4 %, ΔV_30-45min_ = 7.7 ± 2.7 %). (Aii) Spine size changes of unstimulated spines, grouped by distance (< 2.8 μm: ΔV = −14.3 ± 4.7 %; 2.8-6.5 μm: ΔV = 20.3 ± 6.9 %; > 6.5 μm: 11.7 ± 4.0 %). (Aiii) Spine size changes of unstimulated spines, grouped in 2 μm bins. Inhibition of NO production by L-NAME did not affect spine size changes of either stimulated or unstimulated spines. (B) Pr before and 30 min after the induction cl-LTP. (C) Summary of spine size changes. (D) Summary of presynaptic changes (cl-LTP: ΔPr = −0.29 ± 0.04, L-NAME: ΔPr = −0.02 ± 0.06, N = 8, p < 0.01, cPTIO: ΔPr =0.05 ± 0.05). Statistical significance was tested using Kruskal-Wallis H-test and *post hoc* Dunn’s test and comparisons were made with the cl-LTP group. Data points are shown as mean ± SEM.

We next sought to further confirm the functional role of NO in a more physiological experimental paradigm. To do so, we devised an experimental scheme using acute brain slices that closely followed our optical setup in slice cultures, and we assessed the plasticity of Schaffer collateral-CA1 synapses within relatively close spatial proximity of stimulated synapses yet at a distance at which basal responses remained independent (**Fig. 6, see also Methods**). Along individual stretches of dendrites identified by an intracellular fluorescent dye (AF488) introduced by the patched CA1 neurone, we positioned two stimulation electrodes (SE) 10 - 40 μm apart in order to increase the probability of stimulating sets of synapses located close to each other. We used a cross-facilitation test to ensure the stimulation of independent Schaffer collateral pathways. We induced LTP in one pathway by pairing presynaptic stimulation with strong postsynaptic depolarisation via current injection, which elicited bursts of action potentials, 60 times at 5 Hz, a robust protocol used previously to induce presynaptic LTP (Padamsey *et al.* 2017). We then examined changes in the unstimulated pathway.

**Fig. 6:**
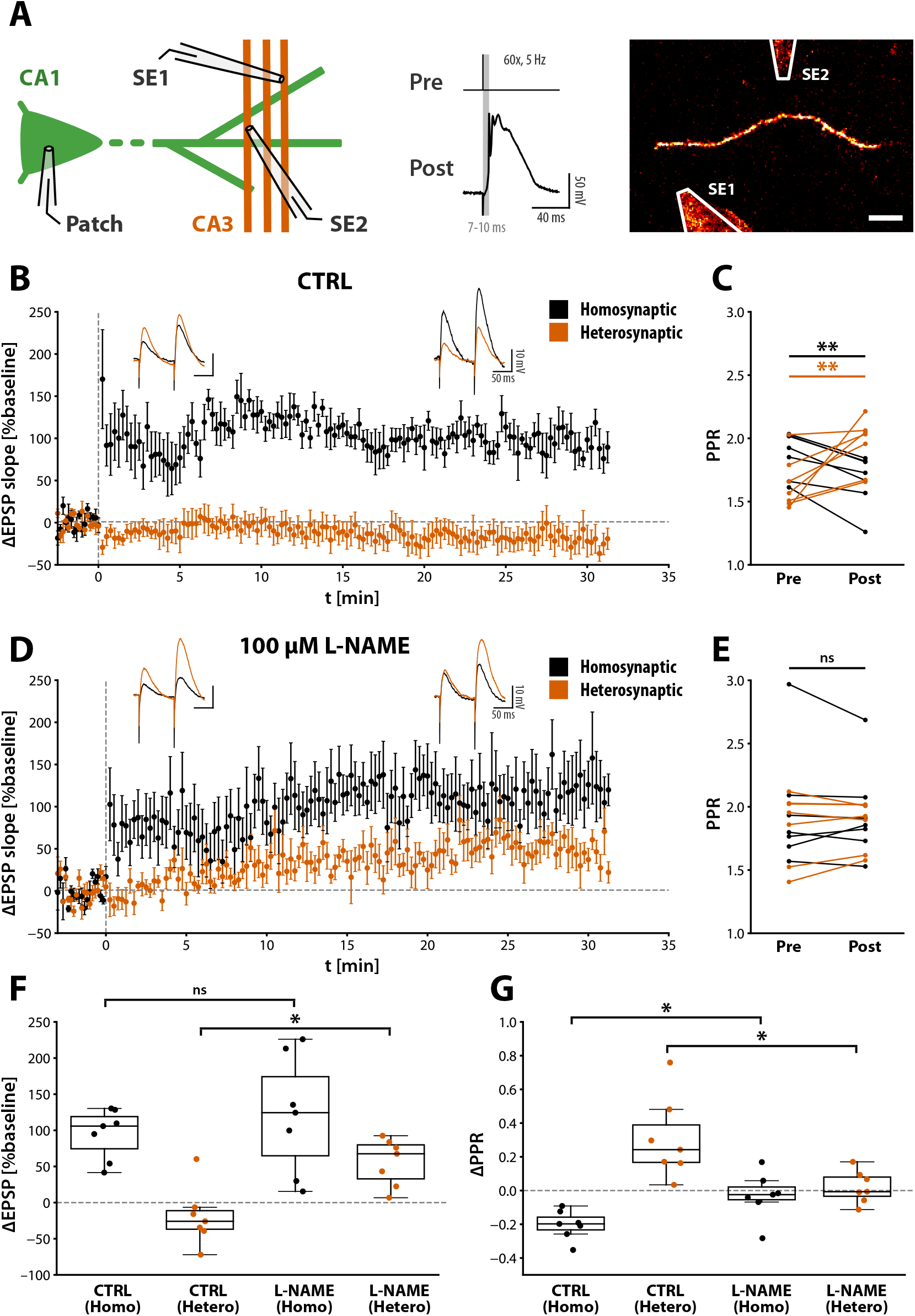
Heterosynaptic LTD in acute hippocampal slices is expressed presynaptically and requires NO signalling. (A) Experimental scheme. CA1 pyramidal neurones in acute hippocampal slices were filled by patching with an electrode containing 200 μM AF488 for 5 min. Two tungsten stimulation electrodes (SE1/SE2) were placed along a segment of dendrite, 10-40 μm apart. Spike timing-dependent LTP was induced in one pathway by pairing presynaptic stimulation with postsynaptic current injection that elicited complex spikes (60 times at 5 Hz). Scale bar: 10 μm. (B) Induction of spike timing-dependent LTP led to long-lasting increase in the EPSP slope of the stimulated pathway (“Homosynaptic”) and decrease in the EPSP slope of the unstimulated pathway (“Heterosynaptic”). Example traces illustrate the average EPSP waveform before LTP induction and during the last 5 min of the recording of the stimulated (black) and unstimulated (red) pathway. (C) PPR before and after the induction of LTP. The stimulated pathway showed a significant decrease of the PPR (Wilcoxon signed-rank test). Conversely, the PPR increased in the unstimulated pathway. (D) Bath application of 100 μM L-NAME abolished heterosynaptic LTD, but did not affect the magnitude of homosynaptic LTP. Spread of LTP was unmasked in the unstimulated pathway. (E) PPR changes of both stimulated and unstimulated pathways were abolished. (F) Summary of EPSP slope changes. (G) Summary of PPR changes. Statistical significance assessed using Mann-Whitney U test. Data points are shown as mean ± SEM.

Induction of LTP led to an increase in the excitatory postsynaptic potential (EPSP) slope of the stimulated pathway (**Fig. 6B**, ΔEPSP = 94.9 ± 13.1 %, N = 7) and a decrease in the paired-pulse ratio (PPR) of EPSP slopes (**Fig. 6C**, ΔPPR = −0.203 ± 0.032), which was stable for at least 30 min, consistent with the previous findings (Padamsey *et al.* 2017). This supported a likely presynaptic expression locus. Conversely, the unstimulated pathway showed mild depression of the EPSP slope with an increase in PPR (ΔEPSP = −19.1 ± 15.4 %, ΔPPR = 0.307 ± 0.092). The heterosynaptic PPR changes were strongly correlated with the fractional change of the EPSP slope (**Fig. S3,** r = −0.842, p < 0.05), which suggested that electrically induced heterosynaptic plasticity was expressed as a decrease in release probability, a finding that was in agreement with heterosynaptic presynaptic weakening that accompanied cl-LTP in slice cultures. In contrast, PPR changes of the homosynaptic pathway were not correlated with the EPSP slope change (r = 0.373, n.s.), which suggested the occurrence of a postsynaptic change that was concomitant to the expression of presynaptic plasticity. Next, we inhibited NO synthase via bath application of 100 μM L-NAME. As observed previously (Padamsey *et al.* 2017), this abolished the homosynaptic presynaptic LTP (**Fig. 6D,E**, ΔPPR = −0.029 ± 0.052, N = 7, p < 0.05). Furthermore, the heterosynaptic increase in PPR was also completely abolished and a stable increase of the EPSP slope was unmasked (ΔEPSP = 56.0 ± 12.3 %, ΔPPR = 0.020 ± 0.036, p < 0.05). NO signalling therefore orchestrates both potentiation and depression of presynaptic strength in a context-dependent manner, that is, potentiation of release at stimulated synapses and depression of release at neighbouring non-stimulated synapses.

## Discussion

Studies of heterosynaptic plasticity to date have mostly focused on local structural plasticity of spines along dendritic branches (Oh *et al.* 2015; Hayama *et al.* 2013; Wiegert and Oertner 2013; Govindarajan *et al.* 2011; El-Boustani *et al.* 2018; Bian *et al.* 2015; Letellier *et al.* 2019; Harvey and Svoboda 2007). Whereas both compensatory and non-compensatory forms of heterosynaptic changes relative to the direction of plasticity induced at the stimulated spines have been reported, the underlying basis for when, where, and what form of heterosynaptic plasticity is expressed have remained unclear. Here, we found that heterosynaptic spine changes that likely represented both compensatory and non-compensatory processes, could be induced at individual synapses by the same cl-LTP stimulus although with different spatial profiles along the dendritic segment. Heterosynaptic LTD was expressed at spines close to the stimulated synapses whereas heterosynaptic LTP was observed at spines further away, which altogether resulted in a spatial profile that resembled a Mexican hat wavelet. What might constitute the emergence of this spatial profile? The most parsimonious explanation of our observations with previous findings could involve the differential diffusion properties of signalling molecules such as those triggered by calcineurin and CaMKII and their interactions with the signalling network in the neighbouring spines (**Fig. 7A, B**). Calcineurin, which promotes structural depression of spines (Oh *et al.* 2015), is itself reported to diffuse into the dendrite and to reach neighbouring spines over short distances (Fujii *et al.* 2013). Furthermore, small GTPases such as H-Ras, Rac, or Rho, which are downstream effectors of the potentiating factor CaMKII, can diffuse substantial distances along the dendrite (Harvey and Svoboda 2007; Harvey *et al.* 2008; Hedrick *et al.* 2016) to mediate heterosynaptic metaplastic changes that manifest as a facilitation to subsequent induction of potentiation (Harvey *et al.* 2008; Hedrick *et al.* 2016).Therefore, differential diffusion along the dendrite of calcineurin and active small GTPases could in effect produce the Mexican hat wavelet profile of spine structural plasticity. In such a scenario, our experiments suggest that the effective diffusion of calcineurin must be weaker than that of CaMKII-dependent signalling events, although this possibility remains to be tested.

**Fig. 7:**
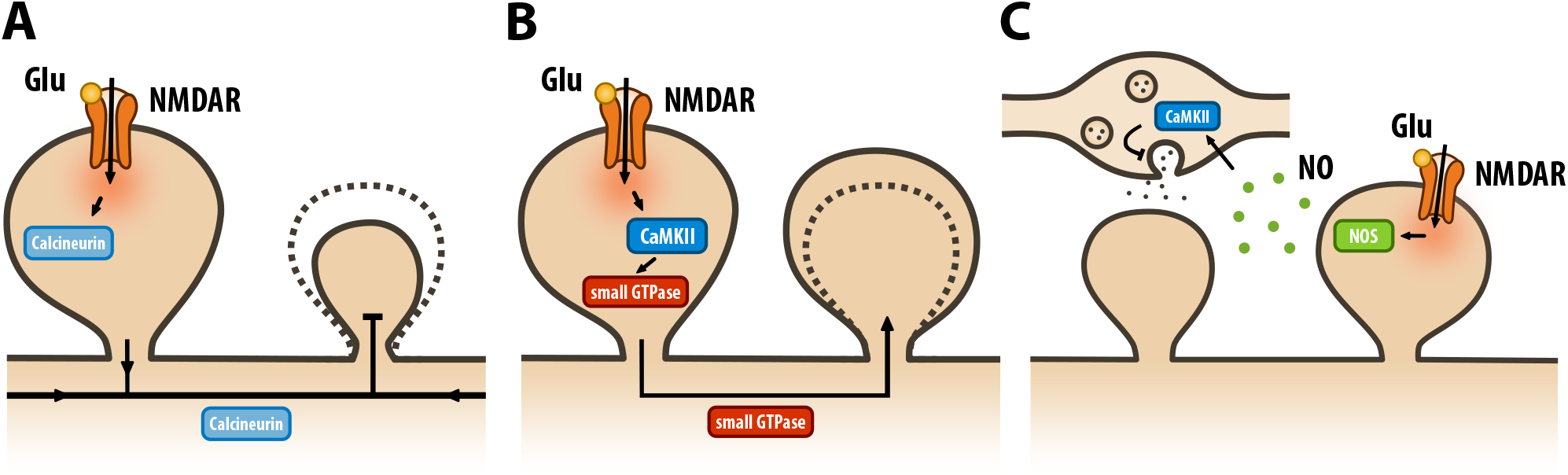
Proposed molecular pathways underlying the local coordination of synaptic strengths. (A) Spatial diffusion of calcineurin, which is activated downstream of NMDARs, leads to heterosynaptic shrinkage of spines. Calcineurin has a high activation threshold but long integration time window and requires the concurrent activation of nearby spines. (B) CaMKII-dependent activation of small GTPases leads to the heterosynaptic enlargement of spines. The diffusion of the GTPases likely extend further than calcineurin to allow for potentiation of distally located spines beyond the reach of the influence of calcineurin that causes spine shrinkage. (C) Activation of NMDARs leads to the synthesis of NO, which acts as a retrograde messenger to decrease Pr at the presynaptic terminal.

The overall spatial pattern of heterosynaptic spine plasticity that we found was mostly consistent with previous reports of heterosynaptic plasticity with some notable differences that were highlighted by the presence of the Mexican hat profile discussed above. The heterosynaptic spine shrinkage that we observed was in accord with the study by Oh and colleagues (2015), where sequential potentiation of spines in close spatial proximity produced shrinkage of unstimulated spines within ~ 3.4 μm of the stimulated cluster in a process requiring the activation of calcineurin but not CaMKII. However, unlike our study, Oh and colleagues did not report an enlargement of spines located further away. Furthermore, several studies of single spine LTP showed heterosynaptic facilitation to subsequent induction of LTP (Harvey and Svoboda 2007; Harvey *et al.* 2008; Hedrick *et al.* 2016). These studies, however, did not observe a shrinkage of nearby spines in contrast to our study and the study by Oh et al. (2015). What might be the cause of these differences in heterosynaptic spine changes accompanying LTP induction at single spines? Previously, CaMKII activation was shown to display a highly supralinear response with respect to stimulation frequency and number (Fujii *et al.* 2013). In contrast, calcineurin activation was sublinear and required a longer integration time window (Fujii *et al.* 2013) from multiple sources (Oh *et al.* 2015), suggesting that calcineurin activation might be strongly thresholded (Oh *et al.* 2015). Based on these findings, we argue that the qualitative differences in heterosynaptic spine dynamics might be explained by the differential activation of calcineurin and CaMKII at stimulated spines, in addition to the difference in the diffusion properties of signalling triggered by these plasticity molecules (see above). In other words, differences in the pattern of stimulation used to trigger LTP could produce distinct pattern of calcineurin and CaMKII activation to bias the plasticity outcome at neighboring synapses. In support of such an idea, the heterosynaptic spine shrinkage that occurred in the absence of potentiation of distal spines was triggered using sequential low frequency (1 Hz) glutamate uncaging of individual spines of groups of synapses (Oh *et al.* 2015). This paradigm involving a slow integration time window between stimulation would favour activation of calcineurin over that of CaMKII, and heterosynaptic LTD would dominate. By contrast, stimulation to elicit LTP that is confined to a single synapse might not reach the activation threshold of calcineurin while favouring the supralinear activation of CaMKII, and in the absence of heterosynaptic LTD, heterosynaptic LTP or metaplasticity for subsequent potentiation would be promoted. In our study, we stimulated groups of synapses quasi-synchronously using glutamate photolysis, which likely facilitated the activation of both CaMKII and calcineurin to sufficient levels. Since stimulation was clustered spatially, calcineurin could locally accumulate and lead to shrinkage of spines close to the stimulated spines. Although yet to be tested, efficient diffusion of CaMKII signalling over those of calcineurin would permit for potentiation of distal spines as discussed above. Notably, when we used electrical stimulation instead to elicit LTP and monitored heterosynaptic plasticity, an experimental setup where we could not precisely control the position of the synapses being sampled, the pairwise distance between stimulated and non-stimulated synapses was likely larger than in the glutamate photolysis experiments, a condition favouring CaMKII activation over calcineurin. While we observed mostly heterosynaptic LTD that was consistent with presynaptic weakening under this condition, blocking this presynaptic depression revealed a component of mild heterosynaptic potentiation (**Fig. 6D-G**). This further supports the idea that calcineurin signalling is spatially confined and heavily thresholded. Taken together, heterosynaptic plasticity is highly sensitive to the ongoing spatiotemporal pattern of synaptic activity, and a better understanding of the coordinated responses of signalling networks to such patterned activity is warranted.

Here we showed a requirement for NO in heterosynaptic presynaptic LTD (**Fig. 7C**), a finding which was somewhat unexpected. Whereas NO has been traditionally studied in the context of presynaptic LTP (e.g. Arancio *et al.* 1996; Stanton *et al.* 2005; Ratnayaka *et al.* 2012; Padamsey *et al.* 2017), several studies have implicated NO in homosynaptic LTD (Zhuo *et al.* 1994; Reyes-Harde *et al.* 1999; Stanton *et al.* 2003; Gage *et al.* 1997).These studies collectively support a putative model for NO-dependent LTD at Schaffer-collateral CA1 synapses that involves the activation of presynaptic internal Ca^2+^ stores via cGMP-dependent protein kinase (PKG), a known downstream effector of NO signalling, along with ADP-ribosyl cyclase, ryanodine receptor, and also requiring the concurrent activation of presynaptic CaMKII (Reyes-Harde *et al.* 1999; Stanton and Gage 1996; Gage *et al.* 1997). The involvement of CaMKII in NO-mediated presynaptic depression is in line with our observation that inhibiting CaMKII abolished presynaptic LTD. Whether postsynaptically generated NO at the stimulated spines diffuses short distances to neighboring presynaptic terminals remains to be confirmed, although the spatially confined heterosynaptic spread of presynaptic LTD (within 4 μm of stimulated cluster) is consistent with the direct involvement of short-lived NO as the extracellularly diffusible messenger.

Previously, a crucial role for NO in presynaptic LTP was demonstrated by the Emptage group (Padamsey *et al.* 2017). Specifically, NO synthesis driven by activation of L-type voltage-gated Ca^2+^ channels (L-VGCCs) coupled to presynaptic activity promoted an increase in Pr.The NO in the current study was unlikely to have originated via the same mechanism due to the fast inactivation of L-VGCCs at depolarised membrane potential used during the induction of cl-LTP. Such differences between the two studies suggests distinct functional roles for NO that depends on the context in which NO is generated. In the previous study, local photolysis of NO at single synapses on its own was not sufficient to induce LTP or LTD in the absence of presynaptic stimulation, whereas after dialysing caged NO into the postsynaptic neurone, photolysing a larger area could elicit LTD (Fig. 5 Suppl. 2 in Padamsey *et al.* 2017). Therefore, induction of heterosynaptic presynaptic LTD might require a critical amount of NO in conjunction with synaptic activity that is dependent on NMDA receptors but not L-VGCCs. In contrast, L-VGCC is needed for NO-dependent presynaptic potentiation.

Lastly, our study did not reveal a strong correlation between pre-and postsynaptic changes.This was especially apparent when we inhibited calcineurin and observed presynaptic LTD in combination with postsynaptic LTP. This disjunction in synaptic plasticity suggests that pre- and postsynaptic plasticity represent independent, parallel processes, which follow distinct learning rules. That the pre- and the postsynaptic sides of the same synapse do not necessarily change their strengths in the same direction may seem counterintuitive based on the overall matching of the pre- and the postsynaptic strengths at basal state across a synapse population (Tokuoka and Goda 2008; Kay *et al.* 2011). Nevertheless, a similar disjunctional plasticity of presynaptic potentiation associated with postsynaptic depression was observed in cultured hippocampal neurones (Xu *et al.* 2013; Letellier *et al.* 2019) and between layer 4 neurones in visual cortical slices (Sáez and Friedlander 2009). Such a mechanism could confer homeostatic stability to synapses although the precise properties of disjunctional plasticity warrants further studies. Notably, the independent regulation of pre- and postsynaptic strengths has recently been explained by the difference in functional impact on the postsynaptic neurone (Costa *et al.* 2017). For future studies on the regulation of synaptic strength, it is therefore crucial to consider both, pre- and postsynaptic strengths for fully interpreting the consequences on neural network activity.

## Materials and methods

### Preparation of organotypic hippocampal slice culture

Organotypic hippocampal slices were prepared as previously reported (Padamsey *et al.* 2017). In brief, hippocampi of postnatal day P6-7 male Wistar rat pups (Harlan UK, Nihon SLC) were isolated and cut into 350 μm-thick transverse slices on a McIlwain tissue chopper (Mickle Laboratory Engineering Co. Ltd. and Cavey Laboratory Engineering Co. Ltd.).The brain was dissected in ice-cold Earle’s Balanced Salt solution (EBSS) containing 35 mM glucose, 20-25 mM HEPES and pH-corrected with 5 mM NaOH to 7.35. Slices were transferred onto cell culture inserts (0.4 μm pore size, Merk Millipore) and placed in a 6-well plate filled with 1 ml/well of culture media containing 50 % Minimum Essential Medium (MEM, Thermo Fisher Scientific), 23 % EBSS, 25 % horse serum (Thermo Fisher Scientific), and 36 mM glucose. Some slices (used for experiments in Fig. 2 and 3) were cultured in media that included 2 % B-27 Supplement (Thermo Fisher Scientific). Slices were maintained at 37°C and 5 % CO_2_ and used for experiments at DIV10-15.

During experiments, slices were constantly perfused (1-2 ml/min) with artificial cerebrospinal fluid (aCSF) containing (in mM) 145 NaCl, 2.5 KCl, 16 NaHCO_3_, 1.2 NaH_2_PO_4_,11 glucose, 2-3 CaCl_2_, and 1-2 MgCl_2_. For experiments with intensive imaging, 1 mM Trolox (Sigma Aldrich) and 0.2 mM ascorbic acid were additionally included. For electrophysiological recordings, 200 nM NBQX (Abcam) was added to prevent strong recurrent excitation due to hyper-connectivity of the slice. The aCSF was bubbled with 95 % O_2_ and 5 % CO_2_ and heated to near-physiological temperature (33-35°C) using an in-line heater (Warner Instruments).

### Preparation of acute hippocampal slices

Acute hippocampal slices were prepared from P14-21 male Wistar rats (Harlan UK). Rats were sacrificed and brains were extracted and immediately submerged in ice-cold dissection media saturated with 95 % O_2_ and 5 % CO_2_. Dissection media consisted of (in mM): 65 sucrose, 85 NaCl, 2.5 KCl, 25 NaHCO_3_, 1.25 NaH_2_PO_4_, 10 glucose, 7 MgCl_2_, and 0.2 CaCl_2_. The cerebellum was manually removed by a coronal cut using a razor blade to create a flat surface, and the brain was glued onto a platform of the vibratome (Microm HM 650V, Thermo Scientific). Brains were additionally stabilised by positioning a block of 2 % agar contacting the dorsal surface of the brain. 400 μm coronal slices were cut and the hemispheres were separated and transferred into a recovery chamber containing standard aCSF (in mM: 120 NaCl, 2.5 KCl, 26 NaHCO_3_,1.2 NaH_2_PO_4_,11 glucose, 1 MgCl_2_, and 2 CaCl_2_), which was bubbled with 95 % O_2_ and 5 % CO_2_. Slices were allowed to recover for 8 min at 37°C and 60 min at room temperature prior to use. Slices were maintained at room temperature up to 4-5 h.

For experiments, slices were transferred to the recording chamber and secured using a “harp” (Warner Instruments). The recording chamber was constantly perfused (2-3 ml/min) with aCSF and 100 μM picrotoxin (Sigma Aldrich) was added to block inhibitory synaptic transmission. The aCSF was bubbled with 95 % O_2_ and 5 % CO_2_ and heated to near-physiological temperature (33-35°C) using a custom made in-line heater.

### 2-photon imaging

CA1 pyramidal cells were bolus-loaded with 0.5 mM AF594 and 1 mM Oregon Green BAPTA-1 (OGB-1) for 45-60 s. OGB-1 was included for optical quantal analysis (see “Optical quantal analysis”). The dyes were given at least 10 min to reach diffusional equilibrium in proximal dendrites and until no obvious change in mean fluorescence intensity over time was observed. After identifying a segment of dendrite with linear geometry that was largely contained within a single optical plain and hence suited for imaging, presynaptic strength was measured via optical quantal analysis. Afterwards, z-stacks (0.5 μm steps) capturing ~ 35 μm of the dendrite centred on the previously measured spine were taken every 5 min. Images were acquired on a Zeiss LSM780 confocal laser scanning microscope with a 63x 1.0 NA objective (Plan-APOCHROMAT, Zeiss) using commercial software provided by Zeiss (Zen 2009, Version 6.0.0.303). Images were taken at 4x zoom, which led to a lateral pixel size of 65.9 nm. Both fluorescent dyes were simultaneously excited using an 800 nm 2-photon laser source (Coherent) and emission was separated using bandpass emission filters. At least three baseline images were taken before applying clustered stimulation to elicit LTP (cl-LTP). Images were then taken at 1, 5, 10, 20, 30, and 45 min after the cl-LTP induction. Experiments showing elevated Ca^2+^ levels or outgrowth of filopodia were discarded as these features correlated to subsequent signs of cell death, such as fragmentation of the dendrite. At the end of the experiment, a z-stack was taken (2 μm steps, 1x zoom), which included the dendrite of interest, the cell body, and the position of the stimulation and puffing electrodes.

### Optical quantal analysis

We performed optical quantal analysis as previously reported (Padamsey *et al.* 2019). In brief, a stimulation electrode (tungsten monopolar in glass pipette) was positioned within 5-10 μm of the dendrite of interest. xt-scans (line scans) covering large fractions of spines were taken at 500 Hz while synapses were stimulated with a paired pulse given at 70 ms inter-pulse interval. The imaging software was synchronized with Clampex software via TTL pulses. Stimulation and the initiation of image acquisition were controlled by Clampex. Responsive spines were detected as time-locked OGB-1 Ca^2+^responses (EPSCaTs). In order to prevent potential confounds of AP failures, which will inflate the estimate of release failures, the stimulation strength was increased until the observed Pr stabilized. For data acquisition, line scans were oriented to capture both the spine of interest and the dendrite, which allowed also the detection of local dendritic spikes or back-propagating APs. Before the induction of cl-LTP, optical quantal analysis was limited to 20-25 trials to prevent photodynamic damage. Optical quantal analysis was repeated 30 min after the induction of LTP and 20-30 trials were taken.The stimulation strength was further increased at the end of the experiment and a paired pulse at 5 ms inter-pulse interval was given to ensure that release events could still be detected.

EPSCaTs were analyzed in ImageJ (Schindelin *et al.* 2012; Schneider *et al.* 2012) and a custom written Python script. Fluorescence signals were background-subtracted, averaged within the spatial dimension, and successful release events were counted manually. All results were cross-validated using an automated analysis script, which detected successful release as an increase in the average Ca^2+^signal within 6 ms after stimulation that exceeded two standard deviations above the mean of the baseline. No qualitative difference was found, however, the automated method was not robust against imaging artefacts or unstable baselines. To distinguish successful release events from dendritic or somatic spikes, the peak intensity was required to be at least 50 % larger in the spine compared to the neighbouring dendrite. Alternatively, the identification of EPSCaTs was aided by the signal onset that preceded dendritic/somatic spikes. Pr was calculated as the ratio between the number of successful release events and number of trials.The sampling error is given by the binomial theorem

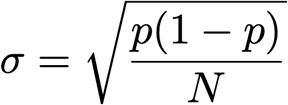

where p is the release probability and N the sample number.

### Glutamate photolysis

Caged glutamate was locally applied during photolysis through a glass pipette coupled to a picospritzer (5-10 psi, Parker Instrumentation). 10 mM MNI-glutamate (Tocris) dissolved in Tyrode’s solution (in mM: 58.44 NaCl, 2.5 KCl, 20 HEPES, 30 glucose, 3 CaCl_2_,2 MgCl_2_, pH-adjusted to 7.2 using 5 mM NaOH) was filtered (0.45 μm pore syringe filter, Merk Millipore) and loaded into a glass pipette (3-4 MΩ).The pipette was positioned ~ 10-20 μm from the dendrite of interest. A 2-photon laser source (720 nm) was used for focal photolysis of glutamate. Photolysis was controlled by a custom written script for the imaging software and was synchronized with the electrophysiology via Clampex. Photolysis consisted of 4 ms pulses, and the laser power was adjusted for each experiment to obtain Ca^2+^transients similar in size compared to EPSCaTs measured in the same cell. For the induction of clustered LTP, laser power was reduced slightly (to around ~ 80%) to prevent overexcitation, which would frequently cause excitotoxicity observed as sustained elevated Ca^2+^ in the dendrite. For quasi-synchronous photolysis, the inter-pulse interval was ~ 2.2 ms.

### Induction of structural LTP

Clustered structural LTP (cl-LTP) was induced by quasi-synchronous glutamate photolysis at 5-7 spines paired with postsynaptic depolarisation, 30 times at 0.5 Hz.The postsynaptic neurone was repatched with internal solution containing 100 μM OGB-1 and 50 μM AF594 to reduce the osmotic pressure of the dye. Photolysis was set up after the GΩ-seal was established and initiated within 10 s of establishing whole-cell mode in order to prevent dialysis. Cells were held at 0 mV in voltage clamp. The average access conductance was Ga = 45.86 ± 2.56 nS. Cells were allowed to reseal after the induction by gently retracting the patch pipette. The maximum distance and mean pairwise distance between stimulated spines did not vary across experimental conditions.

### Analysis of spine structural changes

Spine size was analyzed in ImageJ.The integrated fluorescence of a spine contained in a rectangular regions of interest (ROI) was used as an estimate of spine size. Integrated fluorescence was measured on the z-image slice that resulted in the maximal value. The background fluorescence equivalent to the size of the ROI was subtracted. The re-patch during clustered LTP induction caused a substantial global reduction in the fluorescence intensity. Fluorescence signals were therefore normalized to the mean fluorescence intensity of the underlying dendrite, which was assumed to remain structurally stable. At least three segments of dendrites were averaged to calculate the mean fluorescence intensity of the dendrite. Spine size changes were calculated as fractional changes with respect to the average size prior to cl-LTP induction. Spine size changes at the end of the experiment was defined as the average change observed at 30 and 45 min.

All visible spines along the dendrite of interest were analyzed. Spines that were only partially visible or those located within 2 μm of the photolysis spot were excluded from the analysis. Spines that showed substantial fluctuations (> 25%) during the baseline were also excluded, although these were rare (< 5 % of all spines analyzed). For subgroup analysis of spines, such as distance bins, spine structural changes were averaged within experiments before averaging across experiments.

### Electrophysiological recordings in acute slices

CA1 pyramidal cells located 50-100 μm below the slice surface were targeted for patch clamp recordings. For visualization of the dendritic arbour, 200 μM AlexaFluor 488 (AF488) was included in the internal solution and cells were bolus-loaded for 5 min and patched-off by gently retracting the patch pipette.The fluorescence signal was captured on a Leica DMLFSA confocal microscope with a 63x water-immersion objective (NA = 0.9, HCX APO L 63x/0.9 W U-V-I, Leica) using the Leica Confocal Software (Version 2.61). Neurones were stimulated using two monopolar tungsten electrodes placed directly into the tissue. In order to stimulate sets of synapses in close spatial proximity along individual segments of dendrite, stimulation electrodes were positioned within 10-20 μm of a visually-identified segment of dendrite, 10-40 μm apart from each other. The position of the stimulation electrode could either be visualized via autofluorescence or as shadows in cases where tissue autofluorescence was strong. After positioning, a 10 min wait was included for the stimulation electrodes to reach thermal equilibrium in order to minimize spatial drift. Cells were subsequently re-patched with internal solution containing 200 μM AF488. The probability of successful re-patch was > 90 %. Re-patch of the correct cell was confirmed by fluorescence imaging.

A cross-facilitation test was used to ensure stimulation of two independent pathways. The two pathways were stimulated sequentially with 70 ms inter-pulse interval followed by stimulation in the reverse order, repeated 10 times. A lack of detection of short-term facilitation of the second pulse was used as a criterion for independence of the two pathways. The magnitude of overlap was estimated by comparing the fractional difference in EPSP slope of a given pathway when stimulated as first or second pulse. Significant overlap was rarely observed, which was also supported by the lack of post-tetanic potentiation in the unstimulated pathway.

EPSPs were evoked at 0.06 Hz by 100 μs current pulses (20-80 μA), alternated between the stimulation electrodes. PPR was measured every fifth sweep. A stable baseline was recorded for a maximum of 8 min in order to reduce potential confounds due to dialysis. Experiments showing consistent run-up or run-down were discarded. LTP was induced in one pathway (randomly chosen) by pairing stimulation with strong postsynaptic depolarisation via current injection, 60 times at 5 Hz, as previously reported (Padamsey *et al.* 2017).The unstimulated pathway was kept silent during the induction. EPSPs were recorded for at least 30 min after the induction and PPRs were sampled throughout.

### Statistics

Non-parametric tests were used for all statistical comparisons. Mann-Whitney U test was used for comparisons of independent means; Wilcoxon signed-rank test was used for dependent means. For multiple comparisons, Kruskal-Wallis H test was used followed by *post hoc* Dunn’s test for pairwise comparisons. Sample sizes are indicated in the corresponding sections of the main text. Significance is denoted as follows: *p < 0.05, **p < 0.01.

---

**Fig. S1:**
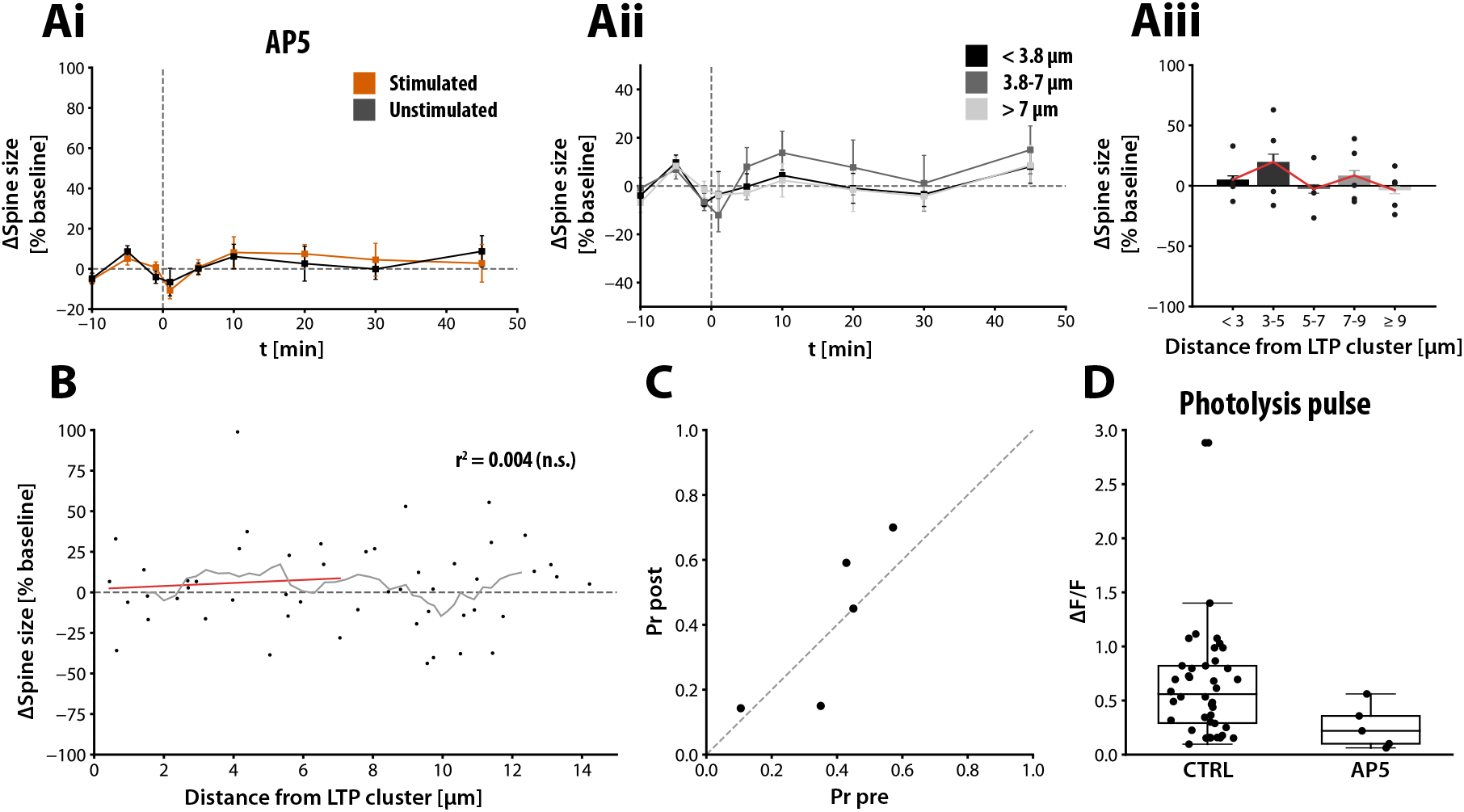
Homo- and heterosynaptic plasticity require activation of NMDARs. Effects of blocking NMDARs by locally applying 500 μM AP5 during cl-LTP induction. (Ai) Spine size changes of stimulated (orange) and unstimulated (grey) spines. cl-LTP was induced at t = 0 min. Both, homo- and heterosynaptic spine size changes were prevented in the presence of AP5 (ΔV_30-45min_ = 4.4 ± 8.1 %; ΔV_30-45min_ = 2.9 ± 5.6 %, respectively, N = 5). (Aii) Spine size changes of unstimulated spines, grouped by distance from the stimulated spine cluster. (Aiii) Spine size changes of unstimulated spines grouped by distance from the stimulated spine cluster in 2 μm bins. (B) Spine size changes of all spines analyzed as function of distance (Pearson r = 0.063, N = 22 spines, n.s.). (C) Pr of unstimulated synapses located near to the stimulated cluster (< 4 μm). No presynaptic LTD was observed in. the presence of AP5. (D) Local application of AP5 decreased spine Ca^2+^transients evoked by uncaging pulses under the baseline condition (CTRL: 0.678 ± 0.092 ΔF/F; AP5: 0.239 ± 0.077).

**Fig. S2:**
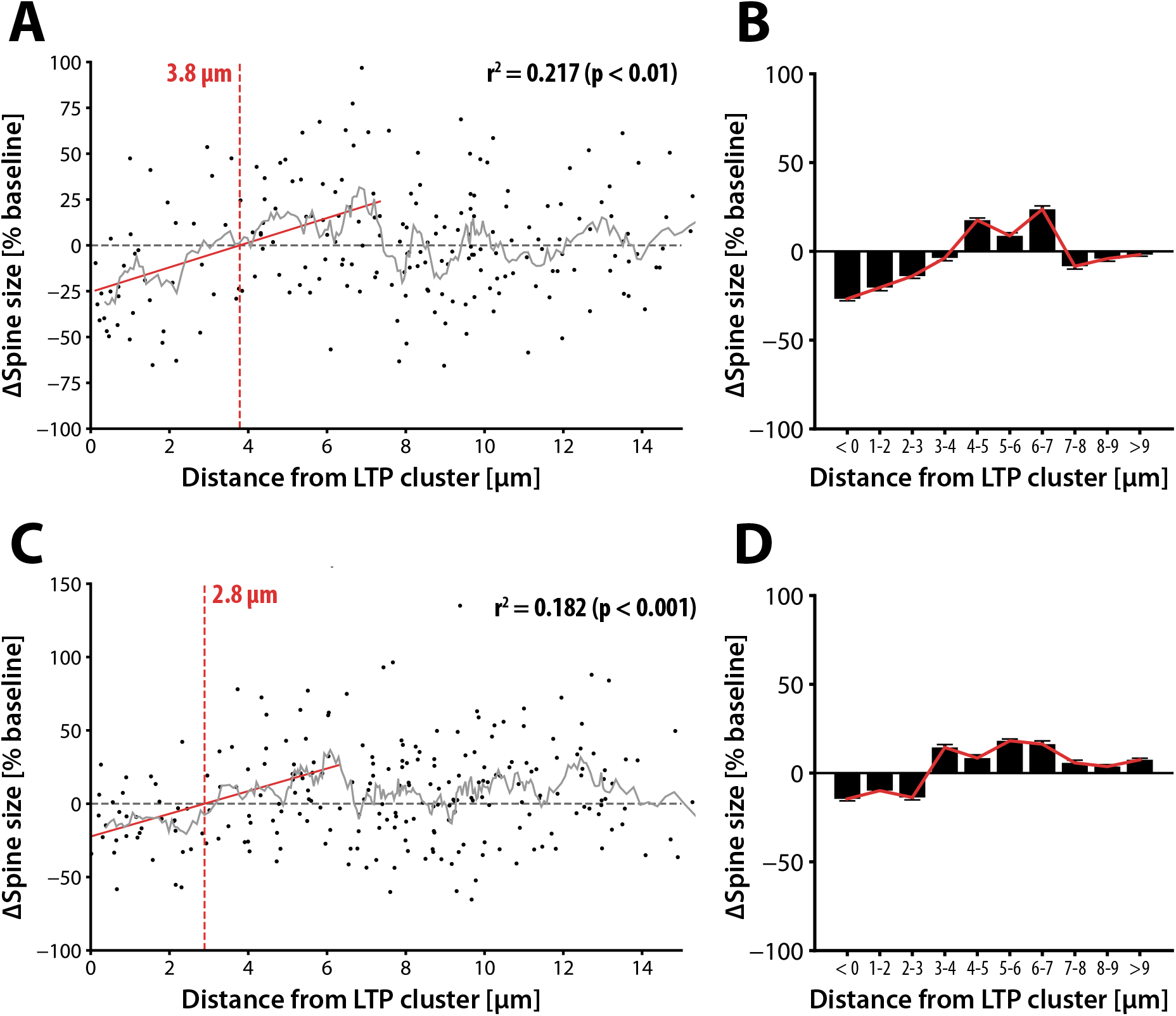
Heterosynaptic spine size change is distance-dependent. (A) Spine size change as a function of distance from the stimulated spine cluster for all unstimulated spines of experiments shown in Fig. 2 (N = 199 spines). The strongest correlation between distance and spine size change was found for spines located within 7 μm (r = 0.465, p < 0.01). The x-axis intercept was 3.8 μm and was used to group spines for further analysis. (B) Spine size changes grouped in 1 μm wide bins from the stimulated spine cluster. Red line highlights the mean spine size change. (C) Distance and spine size change for all unstimulated spines of experiments shown in Fig. 4. The strongest correlation between distance and spine size change was found for spines located within 6.5 μm (r = 0.427, p < 0.01). The x-axis intercept of 2.8 μm was used to group spines for further analysis. (D) Spine size changes grouped in 1 μm wide bins from the stimulated spine cluster. Red line shows the mean spine size change.

**Fig. S3:**
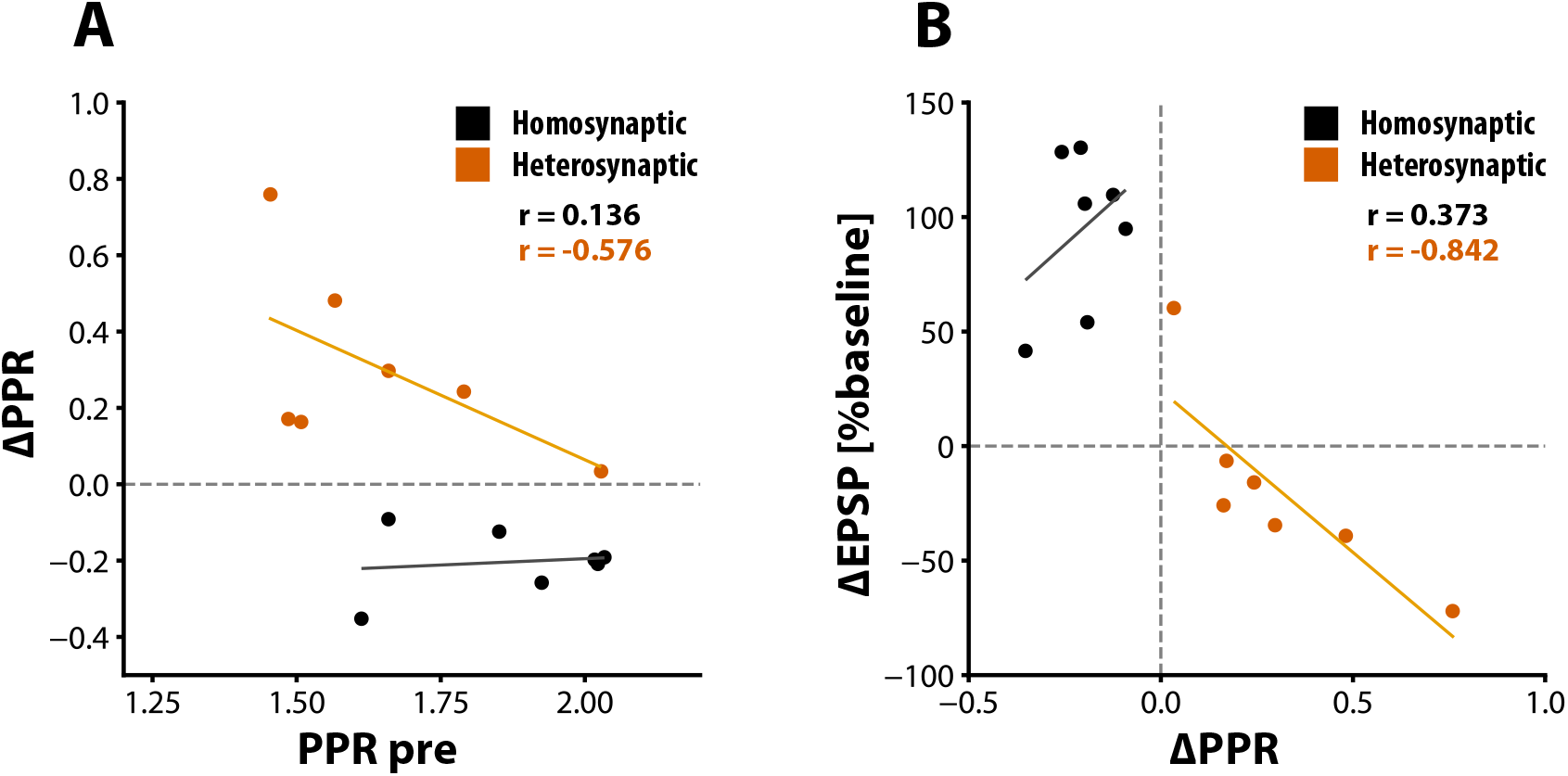
Heterosynaptic plasticity in acute slices is predominantly expressed presynaptically. (A) PPR changes in relation to the initial PPR. PPR changes of the stimulated pathway (black circles) were independent of the initial PPR (Pearson r = 0.136, N = 7, n.s.). Heterosynaptic PPR changes (orange circles) showed moderate correlation, which was however not significant (r = −0.576, N = 7, n.s.). (B) PPR changes in relation to spine size changes. PPR and spine size changes were highly correlated for the unstimulated pathway (orange circles/line: r = −0.842, N = 7, p < 0.05), but not for the stimulated pathway (black circles/line: r = 0.373, N = 7, n.s.).

## Acknowledgements

We would like to thank Tiago Branco, Simon Butt, and Tom Chater for helpful discussion and feedback and Karen Zito for her generous technical guidance. R.T. received the RIKEN-SdV (“Studienstiftung des Deutschen Volkes”) fellowship and was jointly funded by the Clarendon Fund, University of Oxford, and the Medical Research Council UK. The study was also supported by funds from the RIKEN Brain Science Institute, RIKEN Center for Brain Science and JSPS Core-to-Core Program A Advanced Research Networks (Y.G.).

## Author contributions

R.T., Y.G., and N.J.E. conceived the study and designed the experiments. R.T. conducted and analyzed the experiments. The manuscript was drafted by R.T. and all authors edited and approved the final manuscript.

